# Intraoperative Ablation Control Based on Real-time Necrosis Monitoring Feedback: Numerical Evaluation

**DOI:** 10.1101/2023.12.31.573805

**Authors:** Ryo Murakami, Satoshi Mori, Haichong K. Zhang

**Author notes:** Corresponding author: Haichong K. Zhang.

## Abstract

Ablation therapy is a type of minimally invasive treatment, utilized for various organs including the brain, heart, and kidneys. The accuracy of the ablation process is critically important to avoid both insufficient and excessive ablation, which may result in compromised efficacy or complications. The thermal ablation is formulated by two theoretical models: the heat transfer (HT) and necrosis formation (NF) models. In modern medical practices, feed-forward (FF) and temperature feedback (TFB) controls are primarily used as ablation control methodologies. FF involves pre-therapy procedure planning based on previous experiences and theoretical knowledge without monitoring the intraoperative tissue response, hence, it can’t compensate for discrepancies in the assumed HT or NF models. These discrepancies can arise due to individual patient’s tissue characteristic differences and specific environmental conditions. Conversely, TFB control is based on the intraoperative temperature profile. It estimates the resulting heat damage based on the monitored temperature distribution and assumed NF model. Therefore, TFB can make necessary adjustments even if there is an error in the assumed HT model. TFB is thus seen as a more robust control method against modeling errors in the HT model. Still, TFB is limited as it assumes a fixed NF model, irrespective of the patient or the ablation technique used. An ideal solution to these limitations would be to actively monitor heat damage to the tissue during the operation and utilize this data to control ablation. This strategy is defined as necrosis feedback (NFB) in this study. Such real-time necrosis monitoring modalities making NFB possible are emerging, however, there is an absence of a generalized study that discusses the integration and quantifies the significance of the real-time necrosis monitor techniques for ablation therapy. Such an investigation is expected to clarify the universal principles of how these techniques would improve ablation therapy. In this study, we examine the potential of NFB in suppressing errors associated with the NF model as NFB is theoretically capable of monitoring and suppressing the errors associated with the NF models in its closed control loop. We simulate and compare the performances of TFB and NFB with artificially generated modeling errors using the finite element method (FEM). The results show that NFB provides more accurate ablation control than TFB when NF-oriented errors are applied, indicating NFB’s potential to improve the ablation control accuracy and highlighting the value of the ongoing research to make real-time necrosis monitoring a clinically viable option.

## 1 Introduction

Ablation therapy is recognized as a form of minimally invasive treatment, and its application extends to a broad range of anatomical sites, encompassing the brain, heart, kidney, liver, and bone [1, 2]. One of the key benefits of ablation therapy is its capacity for targeted intervention in regions of the body that would be difficult to approach through conventional surgical procedures. For example, glioblastoma (GBM), one of the most fatal types of brain tumor, is sometimes located in a deep area or in close proximity to its eloquent areas. Traditional open surgical methods, often the first option for brain tumors, may be precluded in such circumstances due to the potential risk of inflicting damage on the normal brain tissue or these crucial regions, and ablation therapy can be an alternative option in these cases [3].

The common modalities for high-temperature induction in ablation therapy include 1) radiofrequency, 2) laser, 3) high-intensity focused ultrasound (HIFU), and 4) microwave. They are selected based on an application [4]. Irrespective of the choice, the objective of the therapy is to generate heat adjacent to a target volume, leading to coagulation necrosis, while concurrently circumventing damage to the surrounding normal tissues. Over-ablation may cause complications, while under-ablation could restrict efficacy such as shortened survival rates and recurrence. In fact, for GBM, it is reported that the completion rate of ablation therapy corresponds to the resulting survival [5]. As another example, in the case of cardiac ablation for atrial fibrillation (AF), a considerable recurrence rate is reported. As indicated by Oka *et al*., the recurrence rate of AF stood at 72.7% subsequent to the final ablation procedure (total number of ablation sessions: 1.4 ± 0.7) [6]. Therefore, the precise management of ablation is critical for both safety and efficacy.

To achieve this objective, a couple of ablation control modalities are employed in the current clinical context. The architecture of each control modality is illustrated as a generalized block diagram in Figure 1. The procedural progression of ablation therapy typically includes the following stages: 1) Identification of the target necrosis volume, 2) conversion of the identified target volume into specific control actions of ablation, 3) control of an ablation device according to the determined control actions, 4) induction of heat transferring within the tissue, and 5) cumulation of heat dosage sufficient to induce the desired necrosis formation.

**Figure 1:**
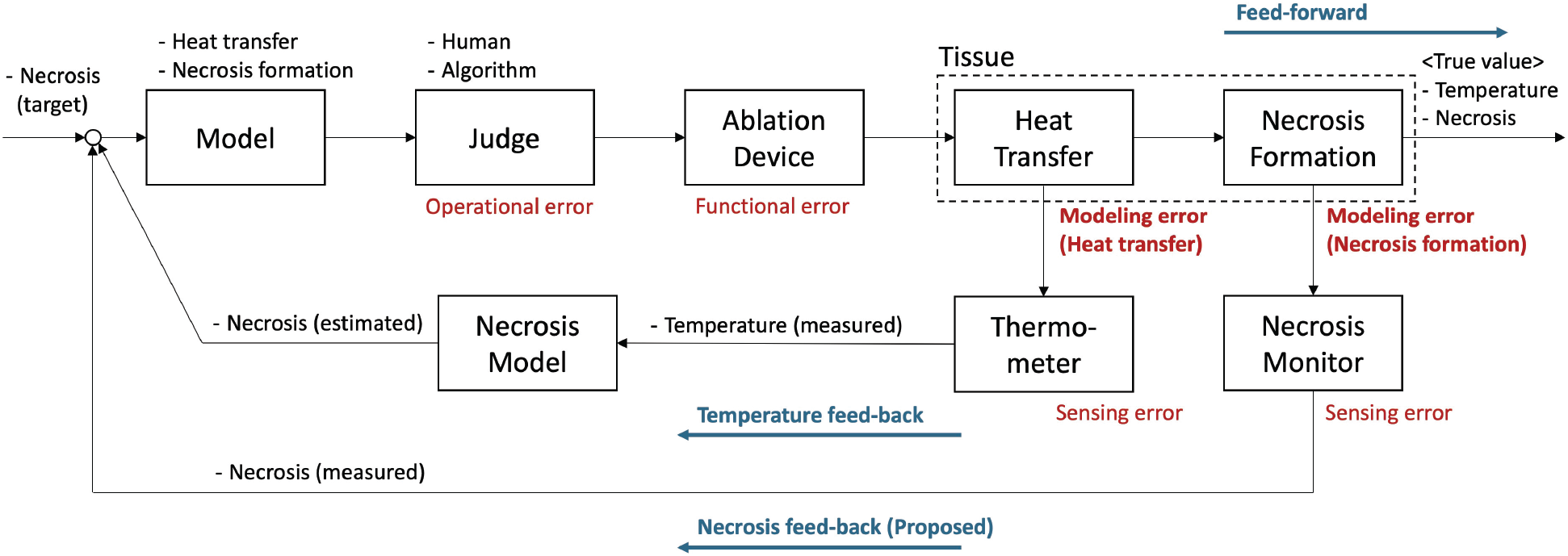
Generalized block diagram of ablation control. Although the temperature feedback loop does not involve the modeling error related to necrosis formation, the proposed necrosis feedback loop does.

A large portion of prevailing ablation therapies employ a feedforward (FF) control mechanism. This approach involves the control sequence pre-defined based on past clinical practice and some theoretical frameworks. When considering cardiac ablation as an example, a conventional control protocol is widely implemented for patients, which includes a prescribed duration of ablation at a specific power for each identified location [6].

Temperature feedback (TFB) is getting more attention as the accessibility of intraoperative thermometers in clinical environments is improved. This methodology permits intraoperative tracking of thermal propagation in tissues during the ablation process, which is then leveraged as feedback data for subsequent control procedures. Since the final goal of ablation therapy is to induce necrosis, the temperature readings obtained via a thermometer are translated into necrosis measurements through a predetermined necrosis model. This type of control is evident in brain ablation procedures where intraoperative magnetic resonance imaging (iMRI) facilitates the monitoring of temperature distribution in the brain during the ablation process [7, 8, 9, 10]. Physicians can subsequently modulate the intensity of laser power, guided by these monitored temperature readings.

Despite the fact that FF and TFB approaches are widely adopted in clinical practice, their control performance is subjected to some constraints. Inside the tissue, two prevailing physical phenomena can be observed: heat transfer and subsequent necrosis formation induced by the accumulated thermal dosage. Though theoretical models representing these physical phenomena have been proposed and investigated [11, 12], it is unrealistic to assume that the parameters and structures of these models apply universally across all patients. Consequently, it is rational to anticipate a certain degree of modeling errors intrinsic to each individual patient and some other environment-related factors.

As depicted in Figure 1, the FF control exerts ablation irrespective of the modeling inaccuracies present in both heat transfer (HT) and necrosis formation (NF) models, thereby leading to the unmitigated propagation of errors into the final ablation outcome. TFB is capable of the intraoperative monitoring of the actual resultant temperature distribution, which involves the HT model’s modeling error. Since closed-loop feedback control is known for its error suppression ability, TFB is expected to suppress the errors originating from the HT model (Appendix A.1.1). While this capability makes TFB more robust against the modeling errors in the HT model compared to FF, TFB is unable to suppress the modeling inaccuracies inherent to the NF model (Appendix A.1.2) because the TFB loop does not involve the actual necrosis including the modeling errors in the NF model.

These aforementioned aspects are intrinsic constraints of FF and TFB when employed in ablation therapy. In other words, those considerations indicate that the feedback control that can directly monitor the necrosis formation would be able to involve the modeling errors associated with both HT and NF models into its closed control loop and suppress these errors (Appendix. A.2), overcoming the constraints of FF and even of TFB.

As the modalities that can visualize necrotic regions, several techniques such as magnetic resonance imaging (MRI), computed tomography (CT), and ultrasound can be listed as the candidates for such a necrosis-based closed-loop control. Previous works investigated the feasibility of these imaging modalities in real-time necrosis monitoring for ablation control [13, 14]. Photoacoustic (PA) imaging is an emerging example for this application, and this modality has shown the potential feasibility in several reports [15, 16, 17] Although the market has yet to widely adopt real-time necrosis monitoring techniques, research proposes several promising solutions, and it is expected that these technologies will soon improve ablation therapy in clinical settings. However, there is an absence of a generalized study that discusses the integration and quantifies the significance of the real-time necrosis monitor techniques for ablation therapy. Such an investigation is expected to clarify the universal principles of how these techniques would improve ablation therapy.

Here, we define necrosis feedback (NFB) control as a general term involving any ablation control with real-time direct necrosis visualization. NFB monitors the necrosis progression intraoperatively and feeds back the information for future ablation control. As discussed above, it is expected that NFB exerts error suppression capability for errors coming from both the HT and NF models.

We evaluate the potential capability of NFB in suppressing errors associated with the NF model. This investigation is conducted in a simulation environment using the finite element method (FEM). We compare the performance of NFB and TFB when artificially generated modeling errors are applied to the NF model. Note that this study does not intentionally presuppose specific modalities for both ablation and sensing, thereby allowing a multitude of potential modalities to extrapolate the findings of this study to their own context.

The structure of this paper is as follows: Section 2 presents the theoretical models used in this study to characterize ablation therapy. It then proceeds to provide details regarding the FEM simulation environment and the parameters employed. The control policy adopted for the purpose of this evaluation, the modeling errors assumed, and evaluation indexes are detailed in Sections 2.3, 2.4, and 2.5, respectively. The implemented simulation is validated by performing both TFB and NFB controls for the HT-oriented errors in Section 3.1. The results of this study are provided in Section 3.2 and the results are discussed in Section 4.

## 2 Materials and Methods

In this section, simulation conditions, the overall flow of the entire investigation, and the optimization of each control method are covered. The global parameters used in this study are summarized in Table 1.

**Table 1:** Global parameters for FEM simulation

| Symbols | Description | Value | Unit |
| --- | --- | --- | --- |
| $T_{\text{duration}}$ | Simulating duration (limitation) | 3600 | Second |
| $T_{\text{step}}$ | Time step | 1 | Second |
| $k$ | Heat conductivity (Human Brain) [11] | 0.51 | W/m/°C |
| $\rho$ | Density (Human Brain) [11] | 1046 | kg/m<sup>3</sup> |
| $c_p$ | Heat capacity (Human Brain) [11] | 3630 | J/kg/°C |
| $T_b$ | Body Temperature (Human) | 37 | °C |
| $T_{a(\text{max})}$ | Maximum ablation temperature | 90 | °C |
| $T_{a(\text{min})}$ | Minimum ablation temperature | 60 | °C |

### 2.1 Biophysical Equations for Simulating Tissue Response

#### 2.1.1 Heat Transfer (HT) Model

In order to simulate the heat transfer inside the tissue, Pennes’ equation is utilized, which is described as shown in Equation 1 [18, 19].

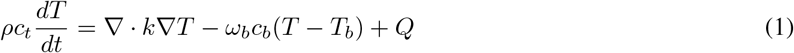

where, *ρ, c*_*t*_, and *k* are the density (kg/m^3^), specific heat capacity (J/kg/°C), and thermal conductivity (W/m/°C) of tissue, respectively. *T* is the temperature at the interested location (°C) and *T*_*b*_ is the blood temperature (°C). *ω*_*b*_ is the blood perfusion rate (kg/m^3^/s), *c*_*b*_ is the specific heat capacity of blood (J/kg/°C), and *Q* represents the heat from other heat sources (W/m^3^). In this study, the terms, *Q* and *ω*_*b*_*c*_*b*_(*T* − *T*_*b*_), are ignored for simplicity.

#### 2.1.2 Necrosis Formation (NF) Model

In general, the process of necrosis formation is a function of both heat and time. There are some proposed NF models representing this process, such as Arrhenius process [20]. In this study, we utilize the Sapareto-Dewey equation as an example [20], which is represented as shown in Equation 2.

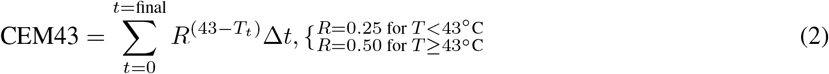

where CEM43 represents cumulative equivalent minutes at 43 °C, Δ*t* is the length of time interval (min), *T* is the averaged temperature during the interval (°C), and *R* is the duration in minutes required to adjust for a temperature fluctuation of 1 °C, either higher or lower than the breakpoint temperature, which is 43 °C in this case [20].

### 2.2 Simulation Environment

#### 2.2.1 FEM environment

Figure 2 (a) represents the geometry of the finite element method-based simulation, which has a rectangular shape with a width of 20 mm and a height of 4 mm. The left side wall of the region corresponds to the body surface, therefore, the horizontal direction can be regarded as the depth in the tissue from the body surface. The ablation device is installed on the left side as a boundary condition, and its temperature (*T*_*a*_) can be modulated between 60 °C and 90 °C. In this study, the performance of each control will be evaluated one-dimensionally; therefore, the horizontal monitoring line is defined at the vertical center of the entire region. Figure 2 (c) is the snapshot of a simulation, where the heat along the left wall propagates in the defined window. The main computation of this simulation including FEM is performed on the MATLAB platform (MATLAB, The MathWorks, Inc., Massachusetts, United States). The mesh grid is made with the condition that the target maximum element edge length is 0.01 mm. In order to perform the simple evaluation of the decision-making capabilities between TFB and NFB, the heat that resides in tissue when the ablation is terminated will be ignored in this study. The temperature distribution is obtained by interpolating the meshes with the resolution of 10 *μ*m. Thus, the length less than 10 *μ*m is discarded when each control error is computed in this study.

**Figure 2:**
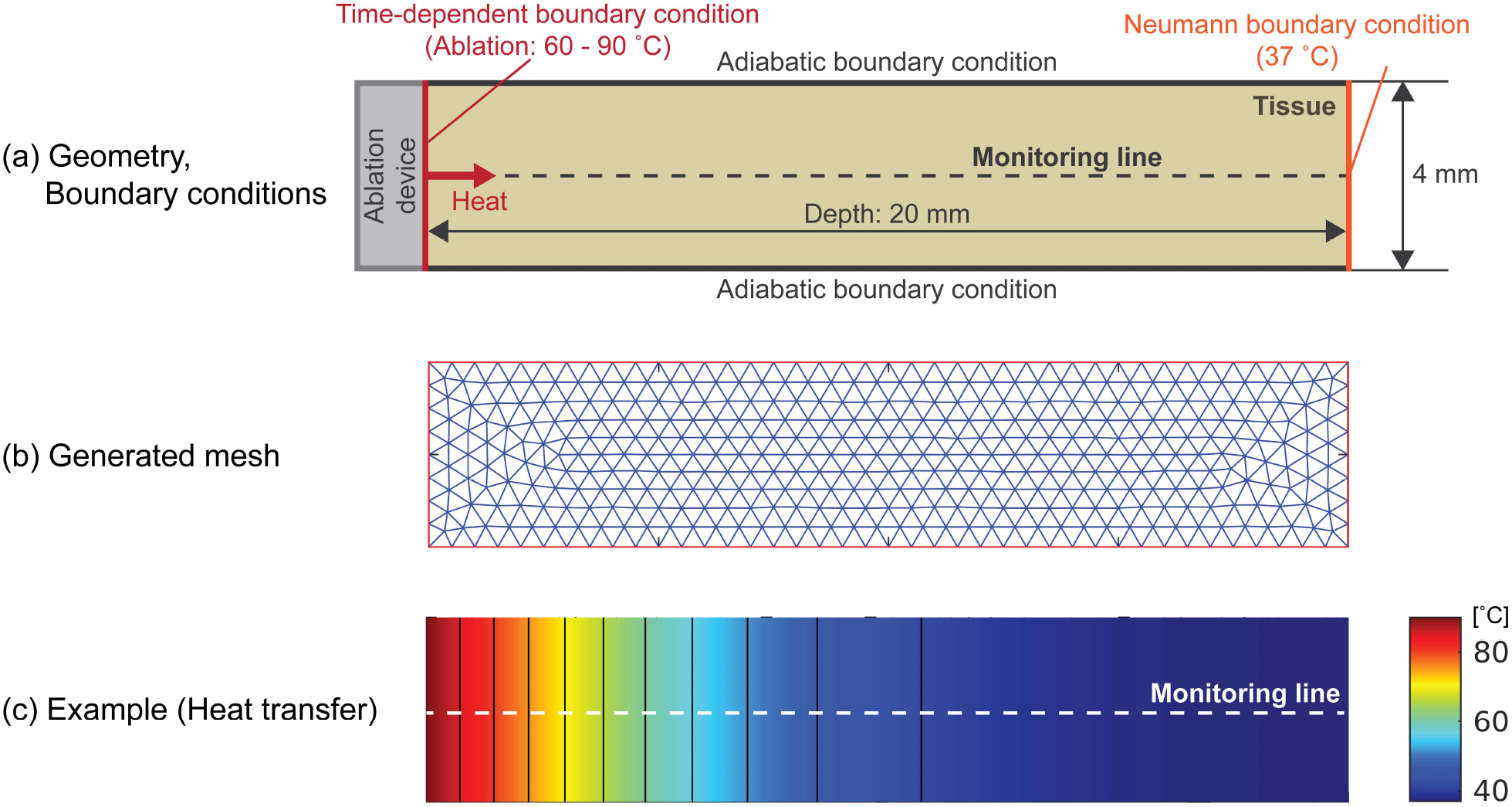
Defined FEM environment; (a) Geometry and boundary conditions, (b) Mesh generated for the thermal simulation, (c) Example of the displayed heat transfer.

### 2.3 Control Policy

To evaluate the contributions of NFB and TFB, the P-control algorithm is implemented as one of the simplest control algorithms. The P controller controls the ablation temperature by computing the multiplication of a pre-defined feedback gain (P gain) and the error value between the targeted necrosis depth and the current necrosis boundary depth. As the current necrosis boundary depth, NFB directly employs the measured actual depth, while TFB estimates the depth based on the assumed NF model and the observed actual temperature distribution. This configuration enables the application of the same P gain for a target depth to both NFB and TFB because the expected types of input and output are the same. This uniformity simplifies the comparative analysis between NFB and TFB. The control cycle is set to 1 sec to be aligned with the simulation time step *T*_step_. The simulation will be terminated when the controller assumes that either the error is small enough or the necrosis boundary exceeds a target depth.

### 2.4 Modeling Errors

To evaluate the control performance of both NFB and TFB when a certain amount of modeling errors are applied to NF and HT models, these modeling errors are generated by randomly determining the parameter values within the fluctuation ranges. The maximum fluctuation range for each error is defined based on the reported property (plugged in the following equations as *P*_lib_) shown in Table 1.

Specifically, the parameter containing an error, *P*_fluc_, is computed as defined in Equation 3.

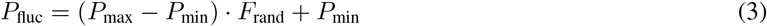

where, *P*_max_ and *P*_min_ are the maximum and minimum values in the fluctuation range, respectively. *F*_rand_ represents the random function returning a random scalar drawn from the uniform distribution in the interval (0,1) (“rand” function provided in MATLAB). In order to provide the identical error conditions to both TFB and NFB, a series of *F*_rand_ was generated in advance and universally used for all the simulations. *P*_max_ and *P*_min_ are defined based on the representative parameter obtained from Table 1, *P*_lib_, and the fluctuation ratio, *r*_fluc_ [%], which ranges from 0 % to 90 % with the 10 %-interval (Equations 4 and 5).

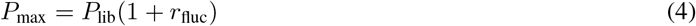

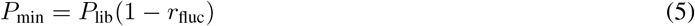

Although there are a variety of parameters that can fluctuate in the both HT and NF models, some of the parameters are selected to simulate the assumed modeling errors. For the HT model, the tissue heat conductivity, *k* is selected as a representative parameter. For the NF model, the CEM threshold value, CEM_th_, that divides the ablated and non-ablated status is selected as this is the parameter determining the robustness of the tissue against heat dose. For CEM_th_, 70 is used as the representing value, *P*_lib_ as it is utilized in other brain ablation studies [21].

### 2.5 Error Index

In order to quantify the performance of each control based on both mean and standard deviation, Error Index is defined as a scalar value combining the mean, *E*_mean_(*i*_*d*_, *i*_*e*_), and standard deviation, *E*_std_(*i*_*d*_, *i*_*e*_) of the control errors as shown in Equation 6. The two values correspond to the errors when the control target depth is *i*_*d*_ mm, and the applied modeling error is defined with the counter, *i*_*e*_. Note that both *E*_mean_(*i*_*d*_, *i*_*e*_) and *E*_std_(*i*_*d*_, *i*_*e*_) are normalized for the absolute maximum among all the combinations of the means and standard deviations. The largest absolute maximum value throughout this study is selected.

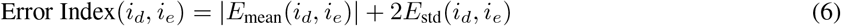

For the comparison of the each control performance with a single metric, Combined Error Index(*i*_*d*_) is also defined (Equation 7) as the combination of Error Index(*i*_*d*_, *i*_*e*_).

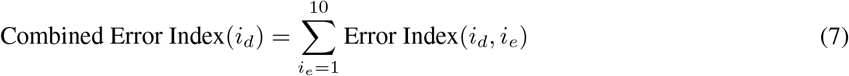

## 3 Results

### 3.1 Validation of Simulation Pipeline

#### 3.1.1 FEM Environment & Biophysical Equations

For the basic validation of the implemented FEM environment and the theoretical models (Equations 1 and 2), trial simulations were performed. One of the trials with these parameters is shown in Figure 4 as time-lapse images. The heat transfer is simulated with Equation 1. CEM is accumulated based on Equation 2, and the CEM profile with respect to depth is visualized in each step of Figure 4. When the CEM at a depth reaches a CEM threshold (e.g., 70), the location will be regarded as ablated. It can be found that the heat is propagated from the left side to the right side. The CEM value at each depth is lifted for time, and the proximal side to the ablator has a higher rate of CEM increase, which corresponds to the definition in Equation 2.

**Figure 3:**
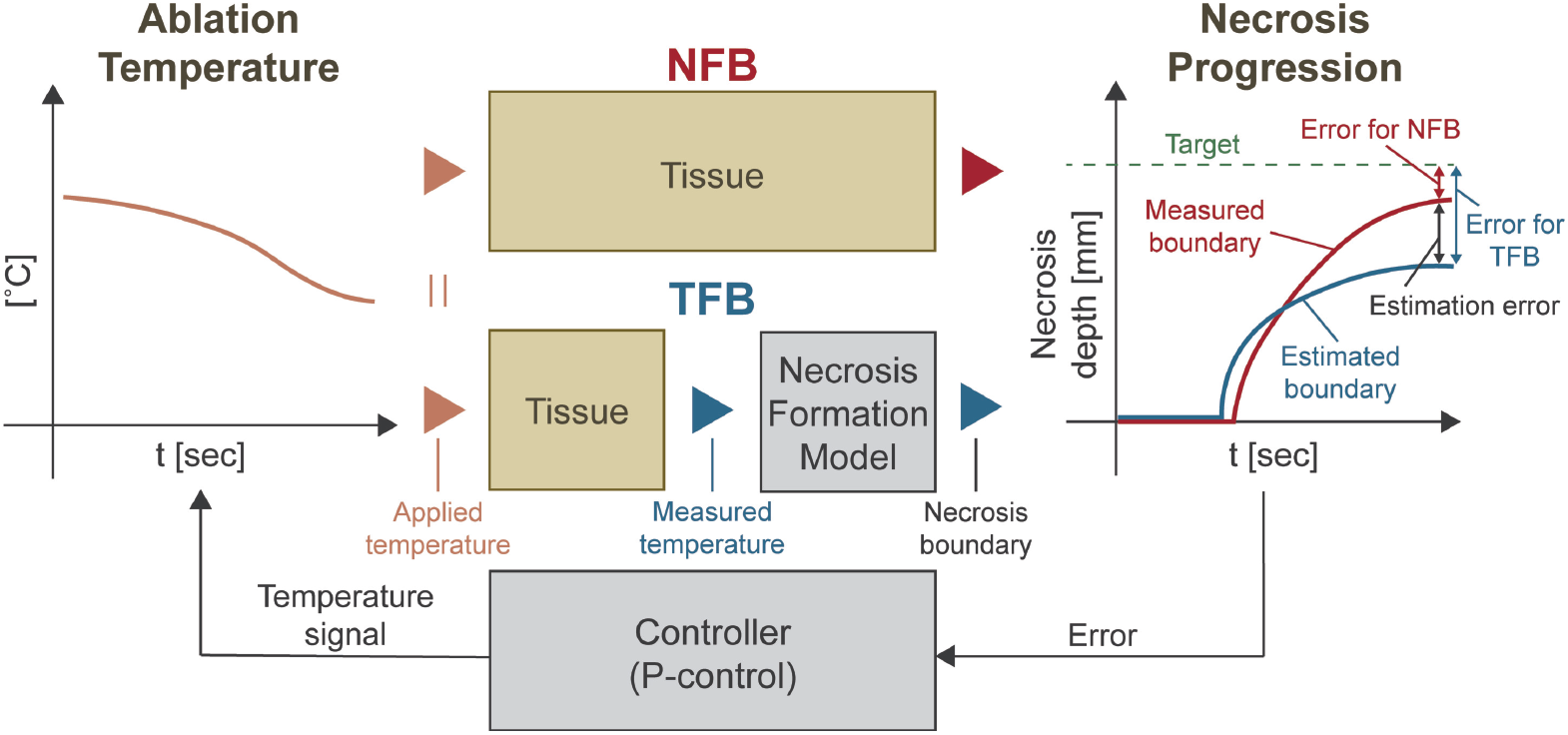
Defined control policy with implemented P controller. The error fed back to the controller is based on the measurement (NFB) or estimation (TFB).

**Figure 4:**
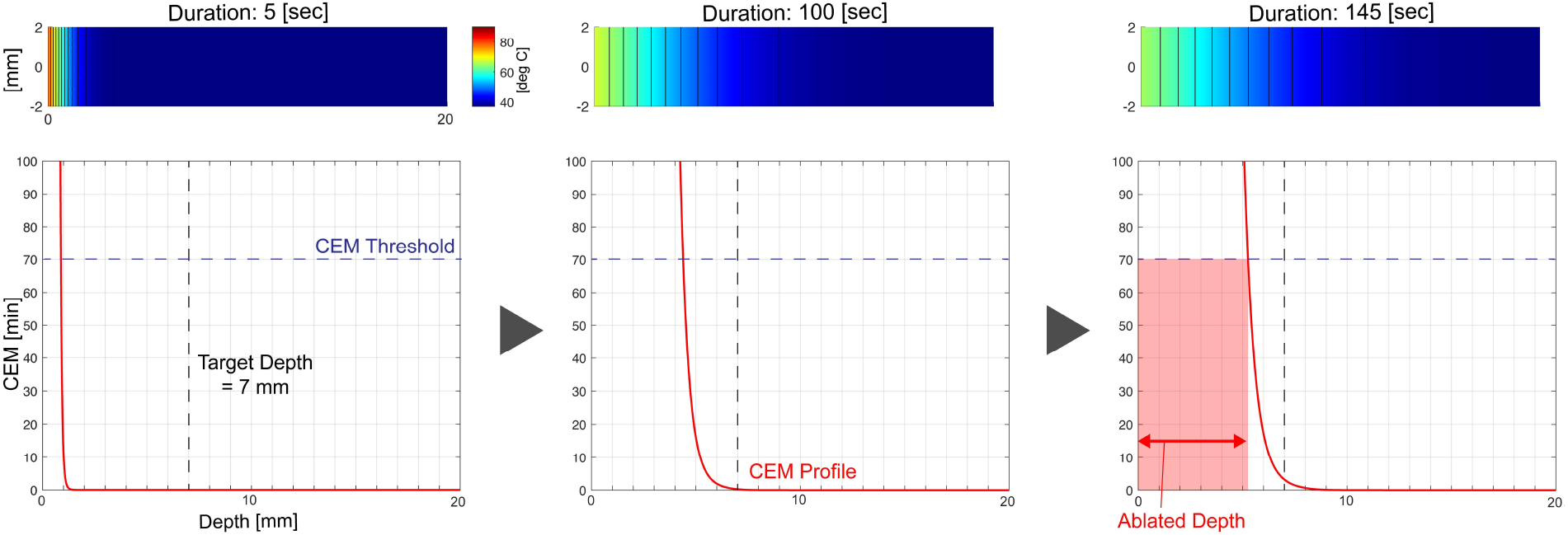
Example of heat transfer and resulting CEM accumulation profile with respect to depth. The heat changes the CEM profile and the region where the CEM profile exceeds the threshold is regarded as ablated.

Based on the observations above, it is confirmed that the implemented FEM with the theoretical models do not contradict their expected behaviors.

#### 3.1.2 Controllers

To validate the implemented NFB and TFB, we observed the control performance of both methods when the errors are applied in the HT model. It is expected that this approach can validate the implemented controllers since NFB and TFB should theoretically provide the same result in error suppression when such errors in the model are applied. As the P-gain, 3000 is universally implemented in this study. The value is determined so that the controller applies the maximum temperature when the control error is 10 mm, which should be the largest number in this simulation. The modeling errors were generated per the protocol described in Section 2.4. For each depth and amount of error, simulation was conducted 100 times in total for both NFB and TFB except for the case in which no modeling error was applied. The mean and standard deviation were computed for each condition and plotted in the graphs in Figure 5. The error is defined as the normalized distance between the target and the necrosis boundary when the ablation was terminated. For the normalization, the maximum values are determined as described in Section 2.5.

**Figure 5:**
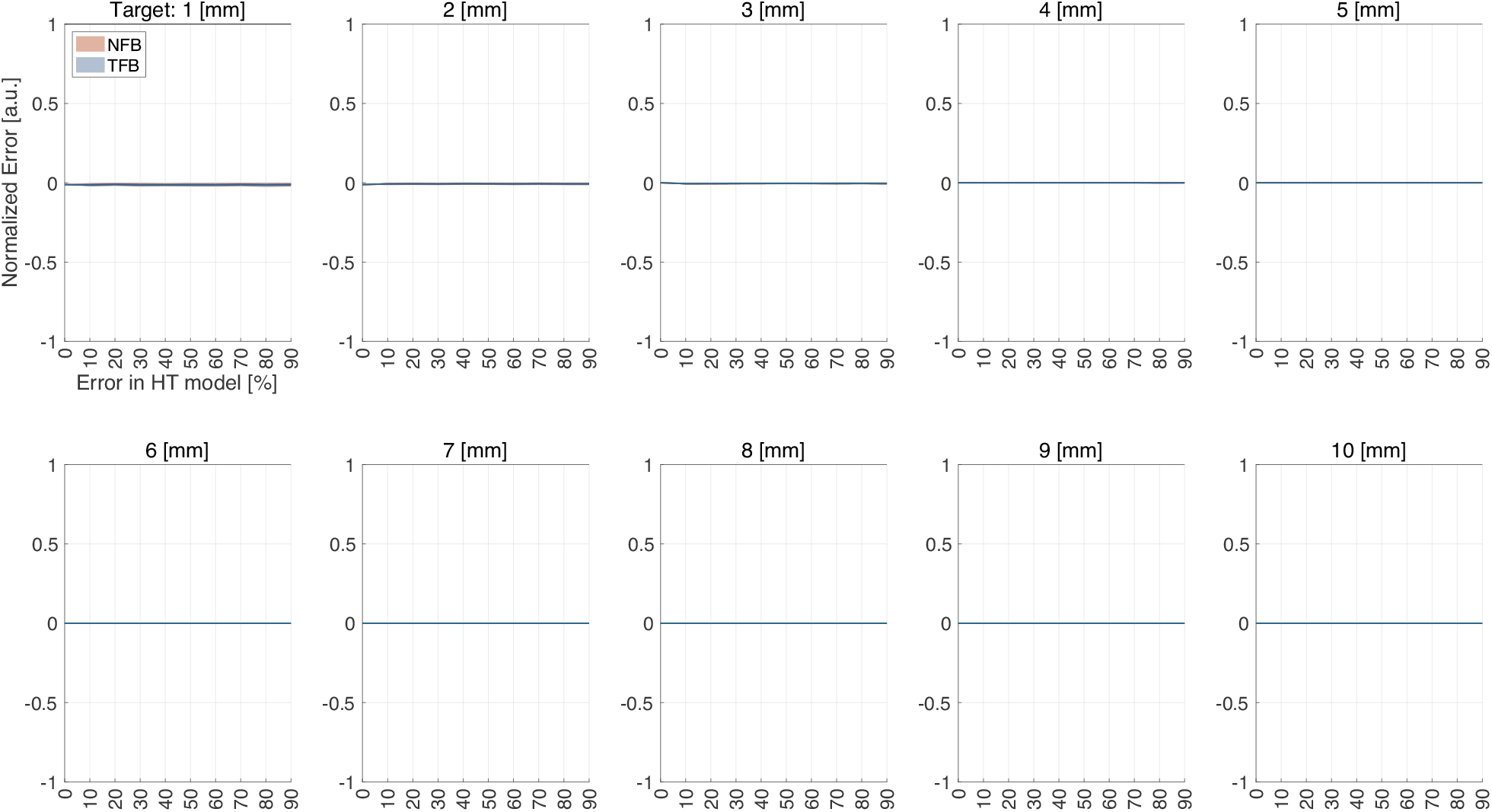
Simulated control errors of both TFB and NFB for each target depth with respect to the applied modeling errors to the HT model.

As shown in Figure 5, both NFB and TFB provide almost no errors when the modeling errors were not applied. In order to investigate how the ablation temperature, *T*_*a*_, is controlled over the ablation process, *T*_*a*_ is plotted as a time series (Figure 6). The 5 mm target is selected as an example. Based on the visual inspection, both NFB and TFB provide the same *T*_*a*_ profiles for all the target depths. It is observed that *T*_*a*_ is first targeted to approximately 74 C° by the controller for all the error amounts in the HT model. This is because the initial distance between the estimated (TFB)/observed (NFB) necrosis depth and the target depth (5 mm) is 5 mm and the same P-gain is used in all the cases. Since the parameter in the HT model fluctuates, it is reasonable that *T*_*a*_ behaves differently because the fluctuation should affect the necrosis progression.

**Figure 6:**
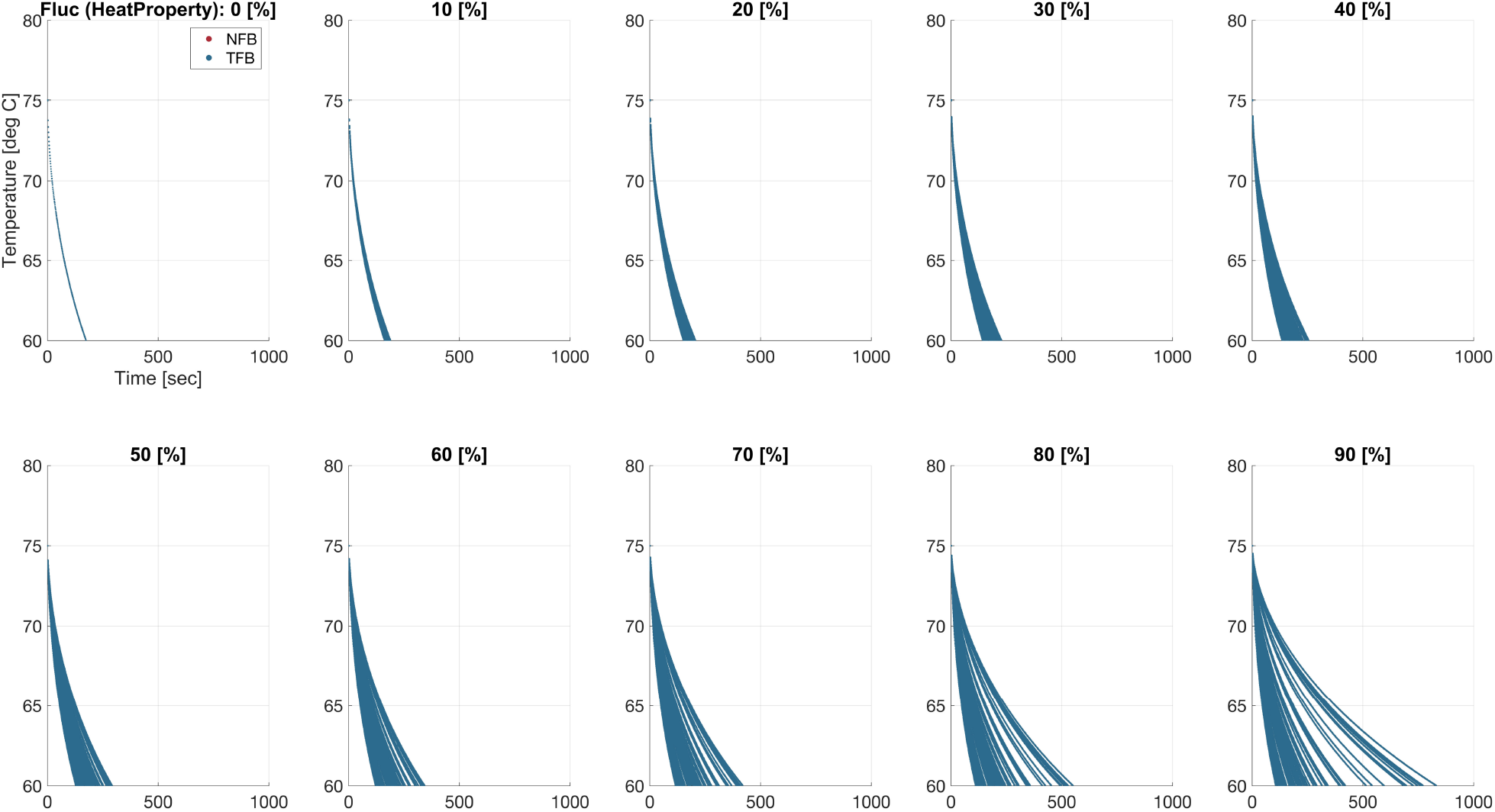
The ablation temperature (*T*_*a*_) reacting to the detected errors in the HT model (Target depth: 5 mm).

To compare the performance of the two control algorithms in a statistical way, the Combined Error Index (Equation 7) was computed for each control, and a two-sided T-test (*α* = 0.05) was performed with the null hypothesis that the Combined Error Indexes of TFB and NFB. The obtained p-value from the T-test was 0.95, indicating the null hypothesis is not rejected. The decomposed Error Indexes (Equation 6) can be visualized as the function of the target depth and error in the HT model (Figure 8). Both TFB and NFB provide almost no errors for each target depth and modeling error. In this case, it is observed that the control performance of TFB and NFB are comparable in each condition. These results do not contradict the expectation that TFB and NFB provide the same control performance when a modeling error exists in the HT model, supporting the validity of the implemented controllers.

**Figure 7:**
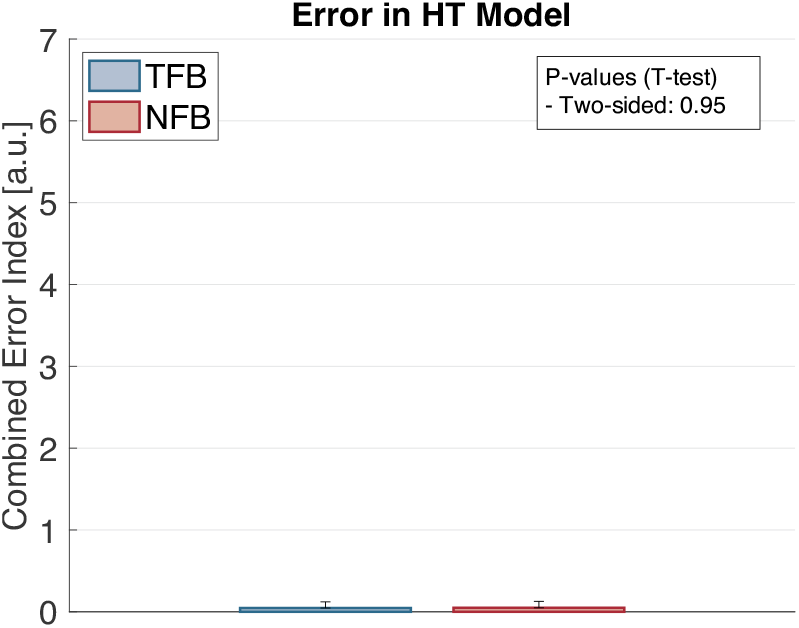
Combined Error Indexes with T-test results for both TFB and NFB when the modeling errors in the HT model are applied.

**Figure 8:**
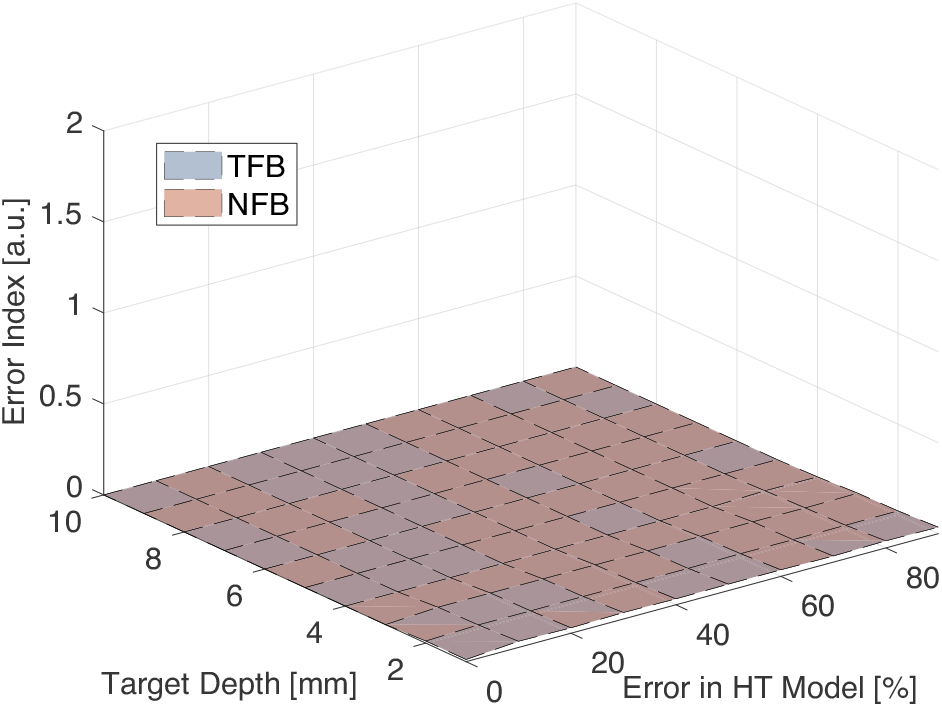
Decomposed Error Indexes as the function of the target depth and error in the HT model.

### 3.2 Evaluation of Necrosis Formation Model-oriented Error Suppression Capability

The purpose of the simulation in this section is to confirm if NFB provides better error suppression capabilities compared to TFB when the NF model has errors, which is the primary research question in this entire paper and expected based on the discussion in Section 1 with Figure 1.

#### 3.2.1 Difference in Control Behavior between TFB and NFB with NF-oriented Error

The artificial errors were applied to the NF model instead of the HT model as performed in Section 3.1.2. With the NF-oriented error, it is observed that TFB and NFB provide different control performances even with an identical modeling error (Figure 9). Error in the NF model is defined as the fluctuation of the CEM threshold, meaning its value can shift from the default value of 70 min, which is assumed as a static value in TFB. In the example shown in Figure 9, the actual CEM threshold of the target tissue is set to 30 min, indicating the error in the NF model is approximately −40 min. For both TFB and NFB, the target depth of ablation-induced necrosis is 5 mm. Although TFB and NFB initially behave in the same way (as exemplified at 5 sec and 60 sec), TFB and NFB terminate their ablation at 174 sec and 147 sec, respectively. In addition to the time, TFB left the evident ablation error (Area colored in gray (Figure 9)). NFB, on the other hand, successfully terminates the ablation when the actual ablation depth is 5 mm.

**Figure 9:**
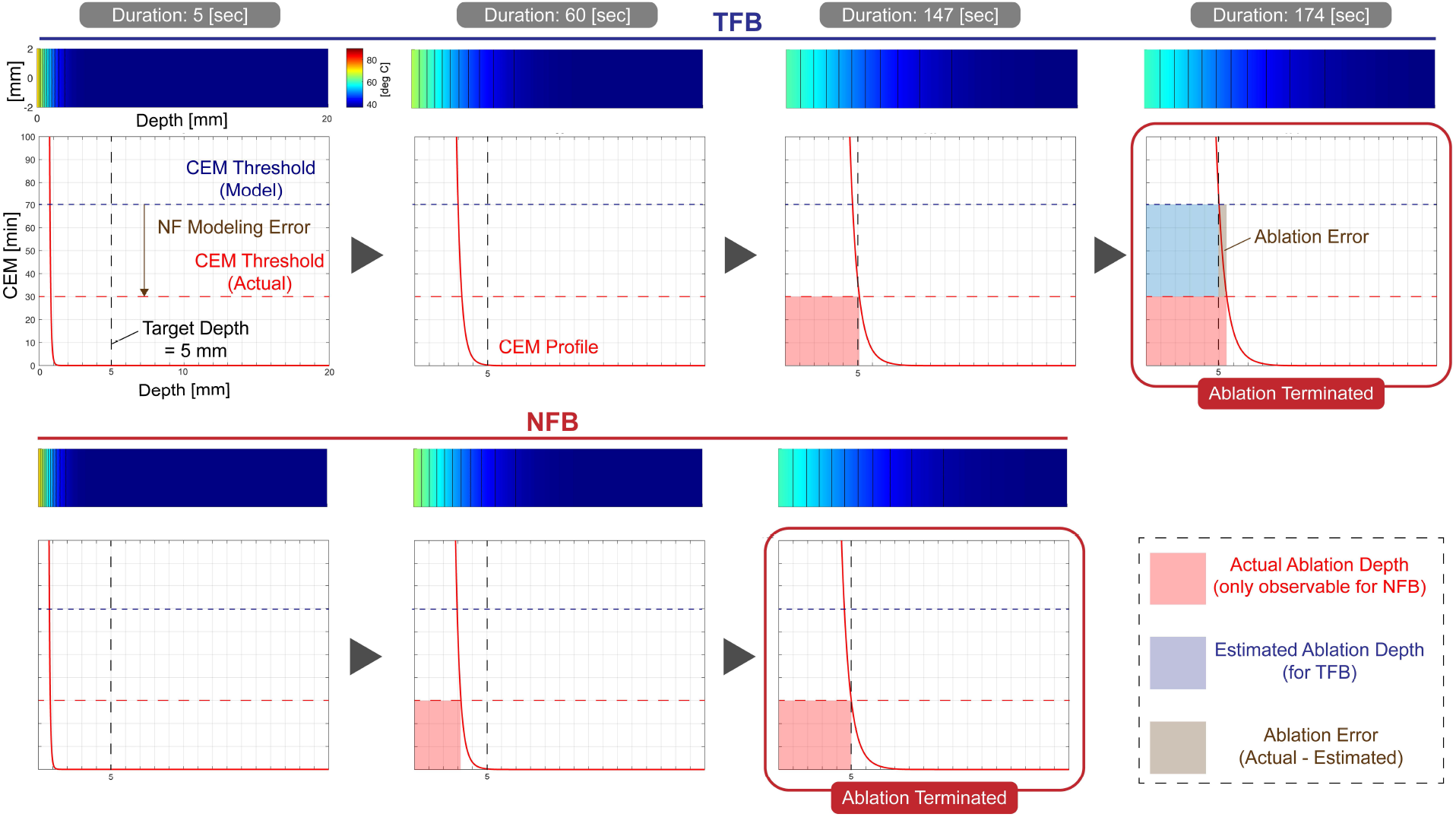
Simulation examples of TFB and NFB with errors in NF model. TFB leaves a certain amount of ablation error because it does not capture the NF modeling error (CEM threshold). NFB terminates the ablation process successfully by considering the modeling error.

As demonstrated in Figure 9, the fixed CEM threshold assumed in TFB can result in the estimation error regarding the necrosis progression, which finally leads to the control error as the TFB control is performed based on the estimation. In fact, TFB terminated the control when the estimated ablation depth (Area colored in blue (Figure 9)) reaches 5 mm, the target depth. On the contrary, since NFB monitors the actual necrosis and feeds back the information to its controller, it properly stops ablation when the actual ablation depth (Area colored in red (Figure 9)) arrives at the target.

#### 3.2.2 Repeated Trials for Statistical Analysis

In order to confirm if the tendency observed in Section 3.2.1 is generally maintained, several magnitudes of the error in the NF model were simulated and control simulation was repeated 100 times for each magnitude like the simulations performed for the HT-oriented error in Section 3.1.2. The same control gain used in Section 3.1.2 was used for these repeated simulations.

As shown in Figure 10, the area representing the standard deviations of TFB is larger than that of NFB in all the cases. Although NFB provides almost no errors at each depth. The ablation temperature modulated by the controller, *T*_*a*_ for the 5-mm depth target starts at around 74 C° for all the error amounts in the NF model (Figure 11). The *T*_*a*_ is the same as the case of the HT model, which is because the errors in neither the NF nor HT model affect the initial control error. Since the same P-gain is shared for all the 5-mm target cases, the resulting control input *T*_*a*_ becomes consistent at the first step of each control. It is observed that *T*_*a*_ for TFB does not fluctuate throughout ablation for all the error amounts in the NF model. This means the P controller for TFB does not capture any errors occurring in the NF model, which is reasonable as the NF-oriented errors are not involved in the closed loop of TFB. In the NFB case, on the other hand, we can observe that the *T*_*a*_ profile fluctuates when the NF-oriented errors are applied, and the amount of fluctuation gets bigger as the amount of error increases. The tendency is explainable because the NF-oriented errors are assumed to happen as the fluctuation of the threshold value for CEM as shown in Figure 9 as an example.

**Figure 10:**
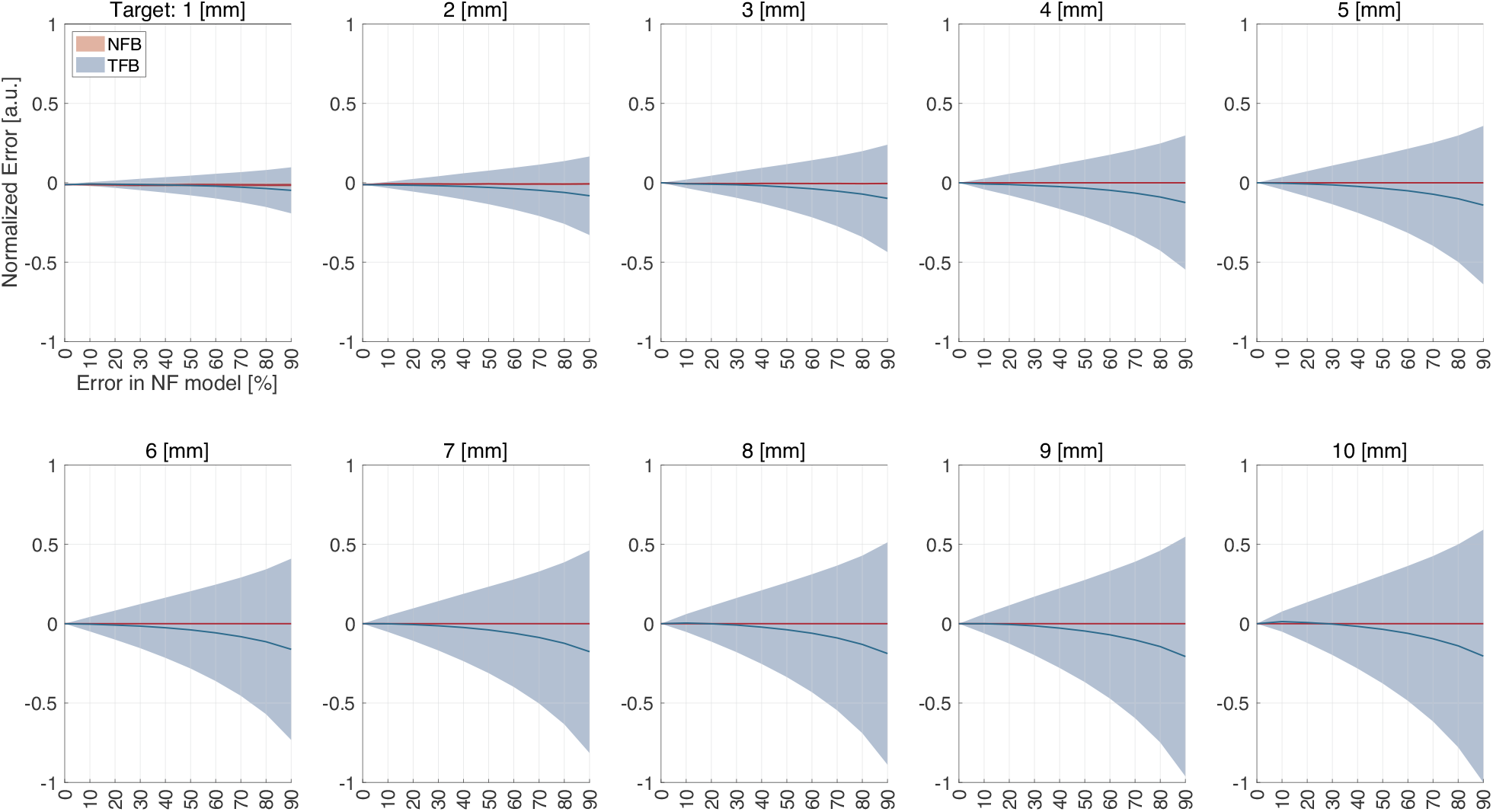
Simulated control errors of both TFB and NFB for each target depth with respect to the applied modeling errors to the NF model.

**Figure 11:**
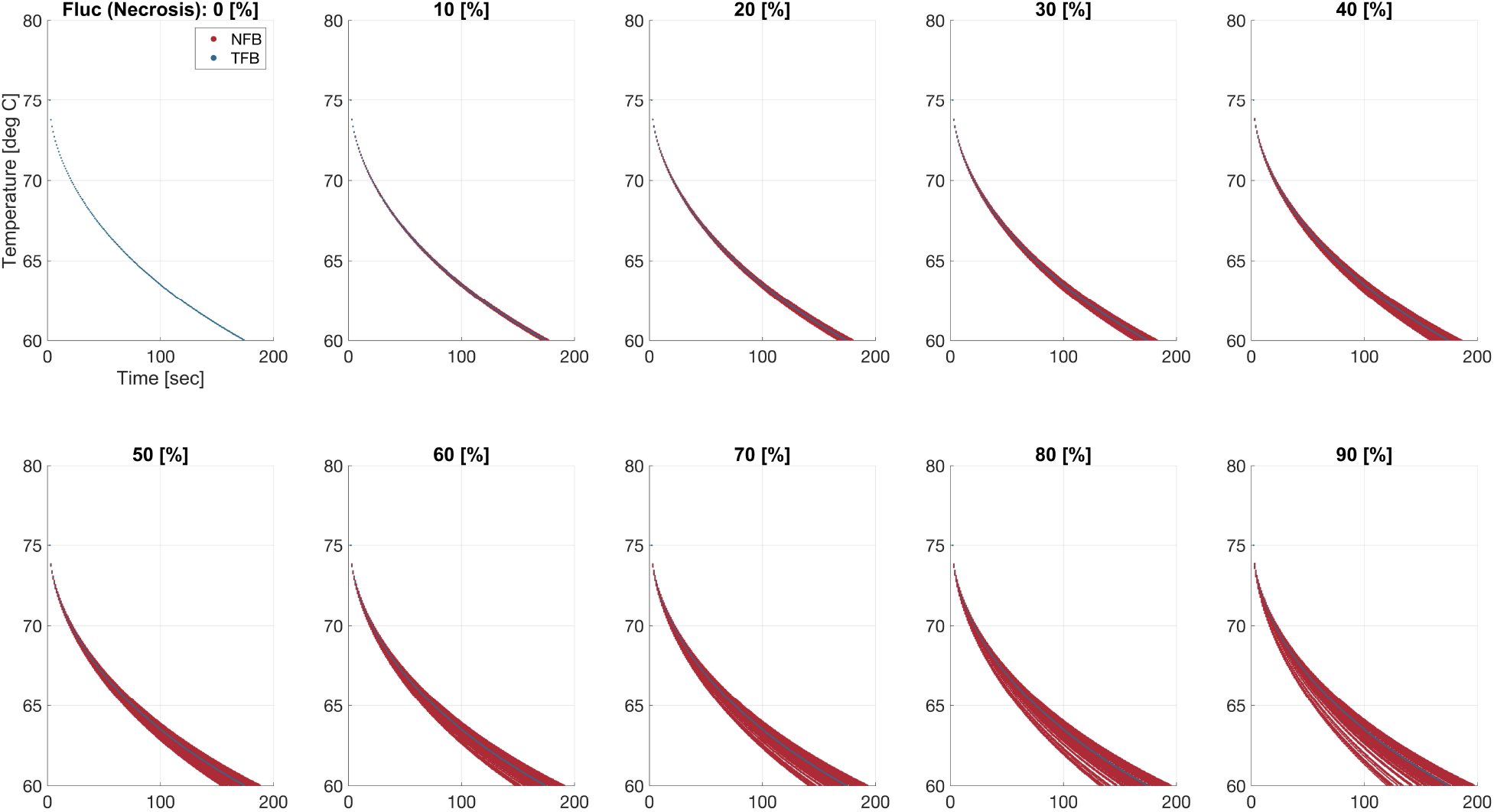
The ablation temperature (*T*_*a*_) reacting to the detected errors in the NF model (Target depth: 5 mm).

For the statistical comparison, the Combined Error Index was computed for each control algorithm, and a one-sided T-test (*α* = 0.05) was performed with the null hypothesis that the Combined Error Indexes of TFB and NFB are equal and the alternative hypothesis that TFB provides the bigger Combined Error Index than NFB (Figure 12). The obtained p-value from the T-test was 0.00, indicating the null hypothesis is rejected. The decomposed Error Indexes are shown in Figure 13. Although NFB provides almost no errors at each depth, the Error Indexes for TFB start to increase once the modeling errors are applied, which corresponds to the statistical analysis done in Figure 12. As a general trend, the increment in the Error Indexes of TFB happens as the errors in the NF model and target depth increase.

**Figure 12:**
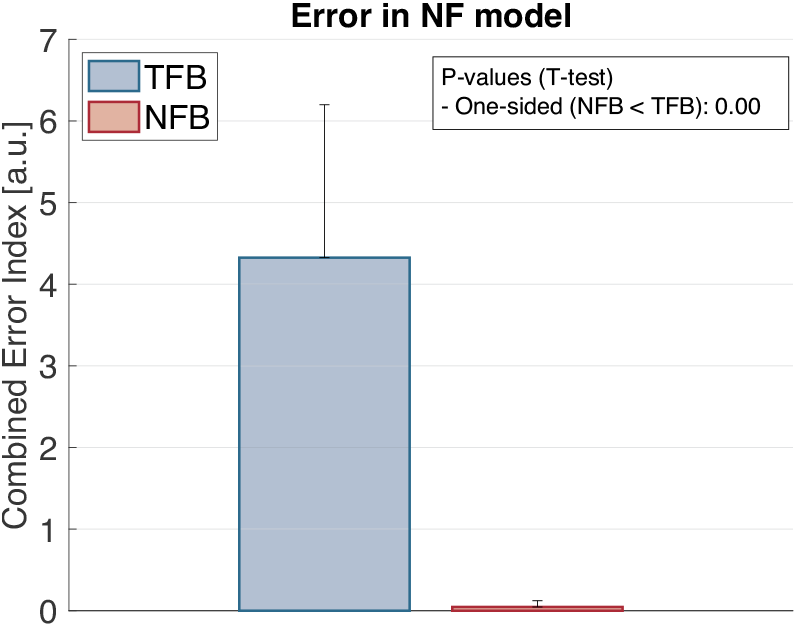
Combined Error Indexes with T-test results for both TFB and NFB when the modeling errors in the NF model are applied.

**Figure 13:**
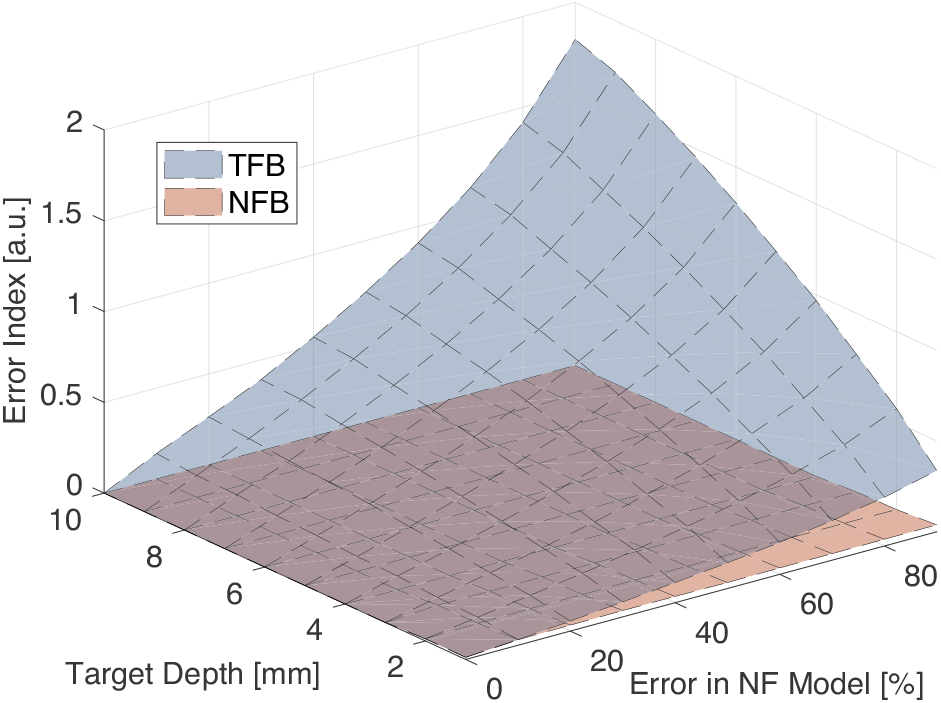
Decomposed Error Indexes as the function of the target depth and error in NF model.

## 4 Discussion

In this study, ablation therapy is simulated in the FEM environment to evaluate how the NFB control realized by real-time necrosis monitoring would improve the ablation control compared to TFB, which is one of the most advanced ablation control modalities in today’s clinical settings. The comparison with the introduced Error Indexes indicates that NFB provides virtually no control errors when the modeling errors occurring in both the NF and HT models are considered while TFB provides a certain amount of errors with the NF-oriented errors. The tendency corresponds to the theoretical analysis of their error suppression capability (Appendix A). The analysis with the simplified model suggests that the closed-loop-oriented error suppression mechanism of TFB does work for the errors in the HT model, but not for the errors in the NF model, and that the suppression of NFB works for the errors in both the HT and NF models. This is simply because the modeling error occurring in the NF model is not fed into the TFB control loop, but is fed into the NFB control loop as described in Figure 1. Given the results shown in Section 3.2, it can be claimed that NFB has the potential to improve the accuracy of ablation control compared to TFB, implying the reinforcement of the importance of the ongoing research aiming at intraoperative necrosis monitoring.

As discussed, one of the key aspects explaining why NFB can suppress the NF-oriented error is the fact that NFB can monitor the actual necrosis progression including errors, and send it back to its controller, but TFB cannot. This can be confirmed from another perspective: the variation of the ablation temperature *T*_*a*_ because it can be the index showing how much each controller tries to be flexible to accommodate the modeling errors. The examples of the *T*_*a*_ variations were presented in Figures 6 and 11 for the target depth of 5 mm. For a quantitative and comprehensive discussion, the *T*_*a*_ variations for all the target depths and amounts of error are visualized (Figures 14a and 14b). Both data in Figures 14a and 14b are normalized with respect to each maximum value among NFB and TFB. Similar to the control errors with the HT-oriented errors (Figure 5), TFB and NFB provide the same *T*_*a*_ variations. In general, it is observed that the variation increases with the increase of the modeling error and the target depth. In the case of the errors in the NF model, the NFB provided less amount of control error compared to TFB (Figure 10), and we assumed this is because only NFB can involve the NF-oriented error in its control loop. As shown in Figure 14b, although the *T*_*a*_ variations in NFB are increasing with respect to the modeling error and the target depth, there are no *T*_*a*_ variations in TFB. These results in Figures 14a and 14b support the claim that both TFB and NFB take into account the HT-oriented error and only NFB is flexible with the NF-oriented error from the *T*_*a*_ variation perspective.

**Figure 14:**
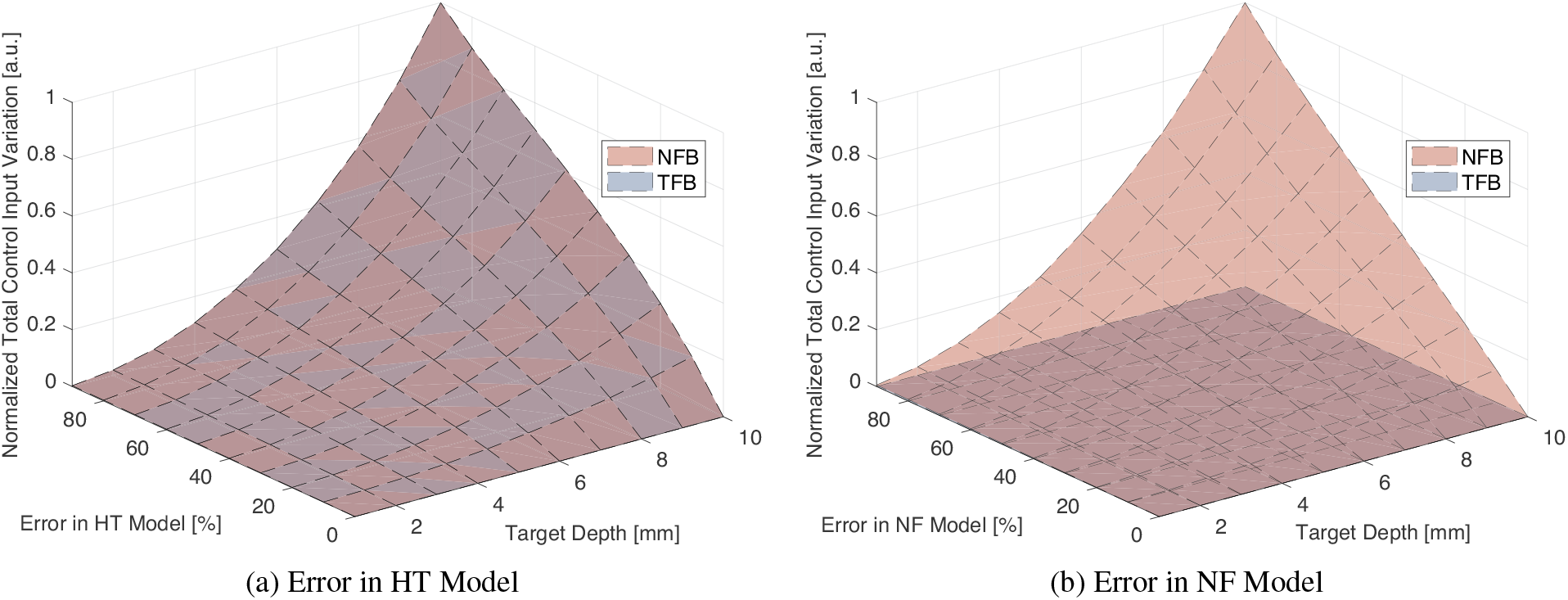
Normalized total variation of the ablation temperature (*T*_*a*_) with respect to the models and modeling errors and the target depths.

Since the simulation in this study was designed with its simplification to evaluate the pure contribution of the assumed real-time necrosis monitoring feedback to ablation therapy, several limitations exist. As defined in Section 2.2.1, the residual heat that should exist when terminating ablation was ignored. Such residual heat would continue to provide heat dose to the tissue for a certain period even after the external ablation is terminated and that would increase the extent of the necrotic region. This effect needs to be considered when we estimate the absolute ablated area, but we decided that it can be ignored in this present study because our primary focus is to see when NFB and TFB decide to stop ablation based on their own monitoring mechanisms. Other types of errors can affect the control performance in the actual control. These errors include an operational error caused by either a human operator or a controller, a functional error occurring within the ablation device, and a sensing error when the heat or necrosis is monitored. Although operational and functional errors are involved in both NFB and TFB closed-loop and the magnitude should be the same for both control algorithms, the sensing error is attributed to each sensor [22, 23]. Therefore, if a sensor for necrosis has greater noise or lower reliability compared to the thermometer used for TFB, the resulting NFB control performance can be worse than TFB. Thus, the specifications of these sensing components need to be considered when an actual sensor is assumed. In this particular study, such sensing error was ignored because the control architecture-oriented difference in performance needs to be evaluated.

These results underscore the need for such a real-time necrosis monitoring modality and also highlight the potential value of the ongoing research being conducted to make real-time necrosis monitoring a clinically viable option.

To study more about the capability of NFB control, we are planning to conduct further simulations and corresponding experiments. These involve the FEM-based simulations considering the residual heat effect and other potential errors such as sensing errors. In addition to these simulation studies, the experiments on tissues comparing the performance of NFB and TFB are being planned.

## 5 Conclusions

In this study, the significance of the real-time necrosis monitoring on ablation control was evaluated by comparing the performance of the NFB control and the TFB control, which is one of the currently most advanced ablation control modalities. The FEM-based simulation suggests that NFB would provide better control performance in terms of the control error at the termination of the ablation when we consider the modeling error occurring in the NF model. Although these results were obtained from the simplified simulation environment, they indicate the potential of real-time necrosis monitoring to contribute to ablation control and imply the importance of ongoing research trying to realize such a real-time necrosis monitoring technology.

## Acknowledgements

This work is supported by grants (CA134675, OD028162) from National Institutes of Health and by Gapontsev Family Collaborative Venture Fund.

## Declarations

During the preparation of this work, the authors used ChatGPT for proofreading in order to enhance the readability and eliminate grammatical errors. After using this tool, the authors reviewed and edited the content and take full responsibility for the content of the publication. The authors have no relevant financial or non-financial interests to disclose.

## A Analytical consideration of error suppression mechanism

This section demonstrates the closed-loop-oriented error suppression mechanism with the simplified block diagram shown in Figure 15. It involves TFB and NFB loops and each physical phenomenon, heat transfer and necrosis formation, is expressed as a scalar value, *M*_*t*_ and *M*_*n*_, respectively. *δ*_*t*_ and *δ*_*n*_ are the modeling errors for each model. P-controller is implemented as assumed in this entire study for both TFB and NFB, and the gains are defined as *K*_CT_ and *K*_CN_.

**Figure 15:**
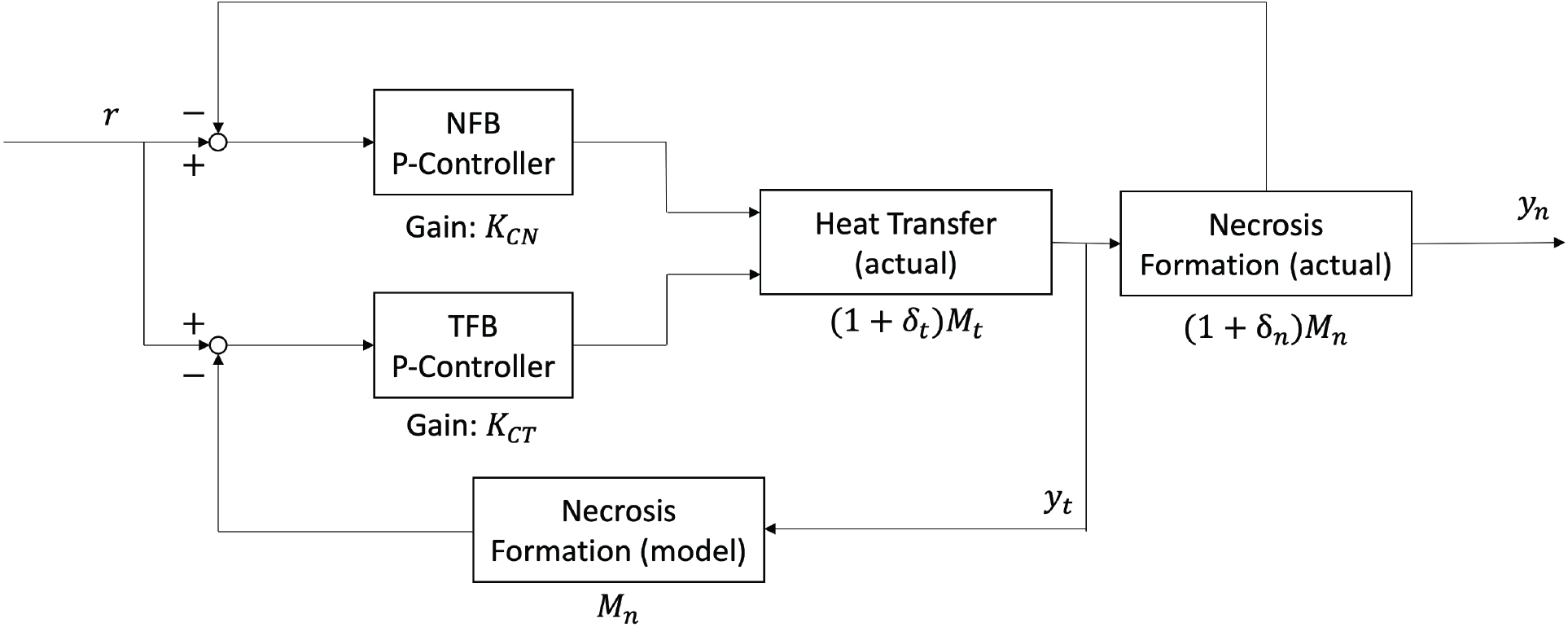
Simplified block diagram.

### A.1 Temperature feedback (TFB)

#### A.1.1 Error in HT model

The target input, *r*, results in the output, *y*_*n*_, based on the structure of the TFB loop. The relationship between *r* and *y*_*n*_ can be represented as shown in Equation 8.

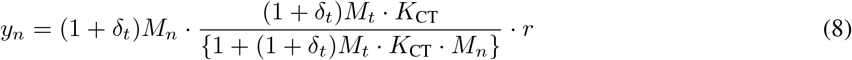

if there is no modeling error in the NF model, then, *δ*_*n*_ = 0. Thus, the following relationship can be obtained (Equation 10)

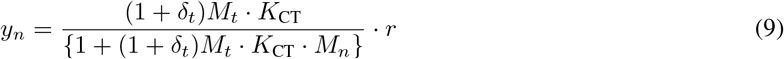

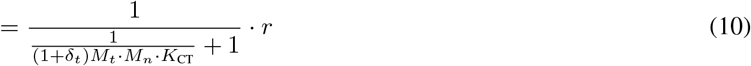

Therefore, the following relationship can be found by increasing the feedback gain, *K*_CT_.

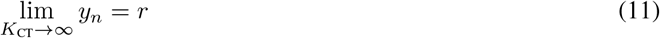

Based on Equation 11, it is confirmed that the modeling error in the HT model, *δ*_*t*_ is suppressed by the closed-loop TFB and *y*_*n*_ is controlled to *r*.

#### A.1.2 Error in NF model

If there is no modeling error in the HT model, then, *δ*_*t*_ = 0. Thus, the following relationship can be obtained (Equation 14).

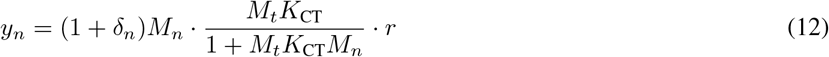

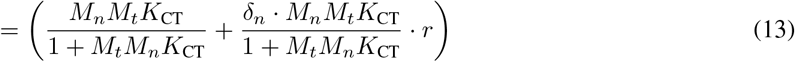

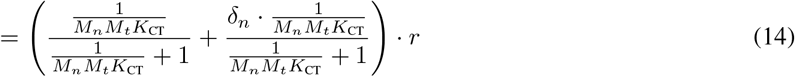

Therefore, the following relationship can be found by increasing the feedback gain, *K*_CT_.

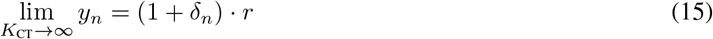

Based on Equation 15, it is confirmed that the modeling error in the NF model, *δ*_*n*_, is not successfully suppressed by the closed-loop TFB control.

### A.2 Necrosis feedback (NFB)

The target input, *r*, results in the output, *y*_*n*_, based on the structure of the necrosis feedback loop. The relationship between *r* and *y*_*n*_ can be represented as shown in Equation 16 and 17.

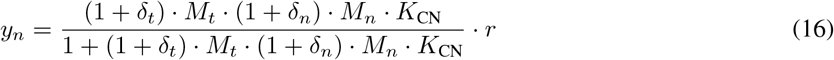

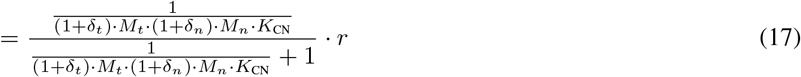

Therefore, the following relationship can be found by increasing the feedback gain, *K*_CN_.

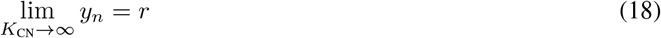

Based on Equation 18, it is confirmed that the modeling errors in both the HT model, *δ*_*t*_, and the NF model, *δ*_*n*_, are successfully suppressed by the closed-loop necrosis feedback control.

